# Static mechanical stretch induces collective alignment of C2C12 myoblasts

**DOI:** 10.1101/2024.10.25.620332

**Authors:** Xuechen Shi, Luyi Feng, Sulin Zhang

**Author notes:** Department of Physiology, University of Pennsylvania, Philadelphia, PA 19104.

## Abstract

Cell alignment is a fundamental process in tissue morphogenesis. While density-dependent collective cell alignment has been widely observed, its underlying mechanisms remain poorly understood. Here, using C2C12 myoblasts, we demonstrate that static uniaxial mechanical stretch induces collective cell alignment in a density-dependent manner: densely populated cultures align robustly, whereas sparse populations do not. We reveal a *biphasic* alignment process, comprising an initial passive phase and a subsequent active phase. The passive phase, driven by substrate deformation, transiently biases cell orientation along the stretch axis regardless of density. In the active phase, initial alignment progressively dissipates in low-density cultures, but is sustained and reinforced in high-density cultures. Supported by coarse-grained agent-based simulations, we propose that self-generated cellular forces facilitate kinetic transitions between orientations, enabling cells to explore orientational states, whereas cell-cell interactions provide a thermodynamic bias that stabilizes the locally aligned state. In dense cultures, strong intercellular interactions promote this stabilization, enabling persistent alignment. In contrast, sparse cultures lack sufficient cell-cell interaction, leading to alignment dissipation. Within this C2C12 system, our findings highlight the cooperative roles of cellular forces and intercellular interactions in orchestrating multicellular ordering, offering new insights into mechanobiology of tissue morphogenesis.

## 1. Introduction

Cell alignment is a ubiquitous and critical process for tissue morphogenesis across diverse cell types. During myogenesis, myoblasts align to fuse into multinucleated myotubes, forming skeletal muscle formation[1, 2]. In blood vessels, endothelial cells align in the direction of the laminar shear stress; disruption of this alignment is associated with inflammatory reactions, endothelial apoptosis, and atherosclerosis[3-5]. During tendon and ligament repair, fibroblasts organize in those tissues by aligning along parallel fibrils, which enhances collagen expression that supports injury healing[6-8]. These examples underscore the functional importance of cell alignment, which has motivated numerous in vitro efforts to replicate this behavior[9-13]. While various molecular and mechanical factors governing cell alignment have been identified[2, 14-20], how multicellular collectives achieve coordinated order remains unclear.

Recent studies highlight cell density as a key regulator of collective cell behaviors. Dense cell colonies exhibit enhanced coordination, enabling cellular responses to subtle stimuli imperceptible to isolated cells. For instance, cell colonies can detect shallow chemical gradient and weak mechanical stiffness gradient, but isolated cells cannot, exhibiting collective chemotaxis[21-25] and durotaxis[26]. Similarly, confluent fibroblasts align on curved substrates while sparsely cultured cells show much weaker alignment, demonstrating collective curvotaxis[27]. On weak topographic patterns, confluent C2C12 myoblasts achieve global alignment, whereas individual cells do not align[28, 29]. Similarly, small human colon carcinoma (HCT-8) colonies display enhanced malignant dispersion[30]. These phenomena converge on a unifying principle: increased cell density amplifies sensitivity to environmental cues, enabling collective responses to weak stimuli. However, the mechanisms by which cell-cell interactions orchestrate collective alignment remain unknown.

Static or quasi-static uniaxial stretch has been shown to align C2C12 myotubes and fibroblasts parallel to the stretch direction[31, 32]. Here, we introduce a new mode of collective cell alignment under static mechanical stretch in C2C12 myoblasts. We identify a *biphasic* alignment phenomenon: an initial passive phase followed by an active, density-dependent phase. In the passive phase, advective substrate coupling causes cells to deform with the stretched membrane, biasing cell orientation; accordingly, cells transiently align along the stretch axis regardless of density. In the subsequent active phase, we propose that self-generated cellular forces, allowing cells to explore their surroundings, enable dynamic transition between orientations, whereas intercellular interactions provide a thermodynamic bias that stabilizes alignment, thus alignment diverges depending on culture density. In sparse cultures, stochastic, cellular forces randomize orientations over time, and insufficient interaction offers no energetic bias to retain order, so alignment dissipates. In dense cultures, strong intercellular neighbor coupling converts the transient deformation-induced bias into persistent order by lowering the energetic cost of co-alignment and promoting cooperative reorientation. Such thermodynamic stabilization outcompetes orientational noise, leading to sustained and progressively increased alignment. At very high packing, however, residual misaligned domains can become kinetically trapped, limiting global perfection. Together, our findings provide biophysical insights into how cellular forces and intercellular interactions cooperatively regulate multicellular organization in C2C12 monolayers.

## 2. Materials and Methods

### 2.1. Cell culture and treatment

Mouse skeletal myoblasts (C2C12) were cultured in Dulbecco’s Modified Eagle Medium (DMEM, Millipore-Sigma) supplemented with 10% (v/v) fetal bovine serum (FBS, R&D Systems) and 1% (v/v) penicillin-streptomycin (R&D Systems). Cells at passage numbers 10-13 were used in the experiments. All cells were incubated at 37°C in an incubator with 5% CO_2_ and 90% humidity. To quantify cell angles, cell nuclei were stained with Hoechst 33342 (Thermo Fisher) at 0.5 μg/ml for 10 minutes.

### 2.2. In vitro stretch application

A stretching device was adapted to uniaxially stretch the cells. The cell stretcher (Electron Microscopy Sciences) comprised a rectangular PDMS membrane that could be mechanically stretched, a membrane chamber for holding the membrane, and a digital distance controller to control the stretch magnitude. This setup allowed for different magnitudes of uniaxial stretch within the membrane. To enable cell attachment, PDMS substrates were incubated with fibronectin (Millipore-Sigma) at 0.05 mg/ml for 30 minutes at 37°C. Upon stretching the cells on the PDMS membrane, mechanical stretch was transmitted to adherent cells through cell-substrate interactions. For angle analysis, the stretch direction was defined to be in the y direction (θ = 90°).

### 2.3. Quantification of cell alignment

Cell orientation angles were determined using the outline-etching method[33, 34], which quantified the angles of cell nuclei to represent cell angles. The degree of cell alignment and directionality to the stretched direction was quantified using an alignment index, a parameter widely used to quantify the alignment level of cells, stress fibers, and ECM fibers[8, 35-37]. The alignment index (AI) was calculated based on individual cell angle (θ_i_):

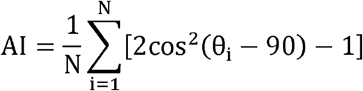

where N is the total number of cells analyzed. If all cells are randomized in their angles, AI = 0; if all cells are aligned to the stretched direction (θ = 90°), AI=1; if all cells are aligned perpendicular to the stretched direction, AI= − 1.

### 2.4. Quantification of deformation ratio

During stretch loading, both the PDMS substrate and the cells were deformed. The deformation, in the y (stretching) or x direction, was defined as the relative change in length of the substrate or cells. The deformation ratio, in the y or x direction, was defined as the ratio between the deformation of a cell and the deformation of the substrate region underneath the cell. A deformation ratio of 1 indicates that a cell adheres to the substrate tightly during stretch; a deformation ratio of 0 indicates that a cell does not change its geometry during stretch. Cell deformation was quantified by tracking the displacement of cell boundaries in bright-field images before and after stretch. To quantify the deformation of the substrate[38], the PDMS membranes were labelled with fluorescent nanoparticles. Specifically, the membranes were treated with (3-Aminopropyl)triethoxysilane (Millipore-Sigma) at 5% (v/v in isopropyl alcohol) for 10 minutes and 0.3% carboxylated fluorescent beads (0.2 μm in diameter, Thermo Fisher) for 10 minutes. The substrates were then passivated with 100 mM Tris base (Millipore-Sigma) for 10 minutes prior to fibronectin coating. Substrate deformation was then measured at the cell scale (>100 μm) from bead-pattern displacements in epifluorescence images, for which the relative error in the deformation ratio is negligible.

### 2.5. Traction force microscopy

To evaluate changes in cell traction forces caused by mechanical stretch, we attached soft polyacrylamide hydrogel to the PDMS membrane as a cell culture substrate based on a previous protocol with modifications[39]. PDMS membranes were treated with (3-Aminopropyl)triethoxysilane at 10% (v/v in isopropyl alcohol) for 30 minutes, washed thoroughly with isopropyl alcohol and water, and dried. They were then incubated with 1% (v/v in water) glutaraldehyde (Millipore-Sigma) for 2 hours, washed, and dried. The membranes were then ready to be coated with polyacrylamide hydrogels.

Polyacrylamide hydrogels embedded with fluorescent beads (0.2 μm in diameter) were prepared on the functionalized PDMS membranes or coverslips to perform traction force microscopy based on previously published methods[30, 39]. After polymerization, hydrogels were functionalized with fibronectin (Millipore-Sigma) by the crosslinker Sulfo-SANPAH (Pierce Chemical)[40, 41]. Traction force was determined by tracing the cell-generated displacement field in the hydrogel by comparing an image pair containing two fluorescent bead configurations in the absence/presence of the cell. To exclude the stretch-generated displacement in the hydrogel from the cell contraction-generated displacement field, the fluorescent image pair was captured with the same stretch magnitude. Initially, an image of the fluorescent beads beneath the cells was taken before applying the stretch (image I). The substrate was then stretched, and cells were incubated for 40 minutes, after which a second image of the fluorescent beads was recorded (image II). Sodium hydroxide solution was then added to detach the cells, and a third image of the beads under stretch was captured (image III). Finally, the stretch was released, and a fourth fluorescent image was taken (image IV). Comparing images I and IV provided the traction force of the cells without stretch, while comparing images II and III provided the traction force after stretch loading.

### 2.6. Laser ablation

Laser ablation was used to kill one cell from a cell pair to study cell-cell interaction and communication. Prior to the experiments, cells were incubated with 2x10^11^/ml gold nanorods overnight. Laser ablation was conducted by irradiating the cell with a laser tweezer module with a single-mode diode laser (150 mW, 1064 nm) in an epi-fluorescence microscope (Nikon, TE2000-U). Detailed experimental procedures for the laser ablation experiments combined with TFM are presented in the Results section.

### 2.7. Agent-based modeling

All agent-based simulations were performed in LAMMPS [42]. Ellipsoidal agents with an aspect ratio of 1:4 were used to represent head-tail-symmetric cells. Cells interact with neighbors via Gay-Berne interactions, incorporating cell-cell adhesion, preferential parallel alignment, and short-range repulsion [43]. The potential between two agents with position vector ***r*_*i*_**, and orientation vector *u*_***i***_ -(cos*ϕ*_*i*_, sin*ϕ*_*i*_), where *ϕ*_i_ is the angle between the long axis and the x-axis, is given by:

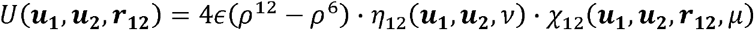

where 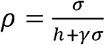 is the shift interaction distance between cells, *h* is the distance of closest approach between agents, is the interaction radius, *η*_12_ and *χ*_12_ are orientation-dependent energy factors, and *∈, γ, v* and *μ* are parameters (Fig. S11) where *∈* sets the overall well depth, *γ* is a shift parameter that sets the location of the potential minimum, and *v* and *μ* are exponents controlling the orientation-dependent prefactors *η*_12_ and *χ*_12_, respectively. The interaction is weakly attractive at long distances and strongly repulsive at short distances. For ellipsoidal agents, the side-by-side parallel alignment is more energetically favorable than other configurations due to the larger contact surface and curvature compatibility.

The two-dimensional simulations were carried out with periodic boundary conditions. The characteristic length, time, and energy were set to be 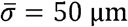, *τ* = 30 min, and 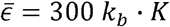 . In a 1 mm by 1 mm simulation box, ellipsoidal cells with a long axis of 100 μm were initially seeded at low density (25 cells) or high density (400 cells), with random, uniform, or domain-based neighboring alignments. The system first underwent energy minimization to resolve instabilities and overlaps, and then multicellular dynamics were simulated using NVE integration with a Langevin thermostat [44] to maintain room temperature for 30 h. The time evolution of agent orientation angles was recorded for further analysis. For the agent-based modeling videos, each timestep corresponds to 3 min.

### 2.8. Statistical analysis

Statistical analysis was performed using Student’s two-tailed t-test. Data are presented as mean ± standard error of the mean (SEM). The significance level of each statistical test is presented with each set of experimental data.

## 3. Results

### 3.1. Mechanical stretch induces collective alignment

To investigate density-dependent cell alignment, we seed C2C12 myoblasts on fibronectin-coated poly(dimethylsiloxane) (PDMS) membranes mounted on a stretch device. Cells attach and spread within four hours of seeding; if allowed to proliferate to confluence over ∼3 days, they do not show apparent alignment (Fig. 1A and Fig. S1A). Prior to stretching, cells are seeded at varying densities (near-confluent high density and sub-confluent low density) and incubated overnight. A uniaxial stretch protocol is then applied: three sequential 15% static stretches (45% cumulative strain) with four-hour intervals, are delivered using a distance controller. Each stretching takes only a few seconds, and the stretch does not alter cell viability (Fig. S2).

**Figure 1.**
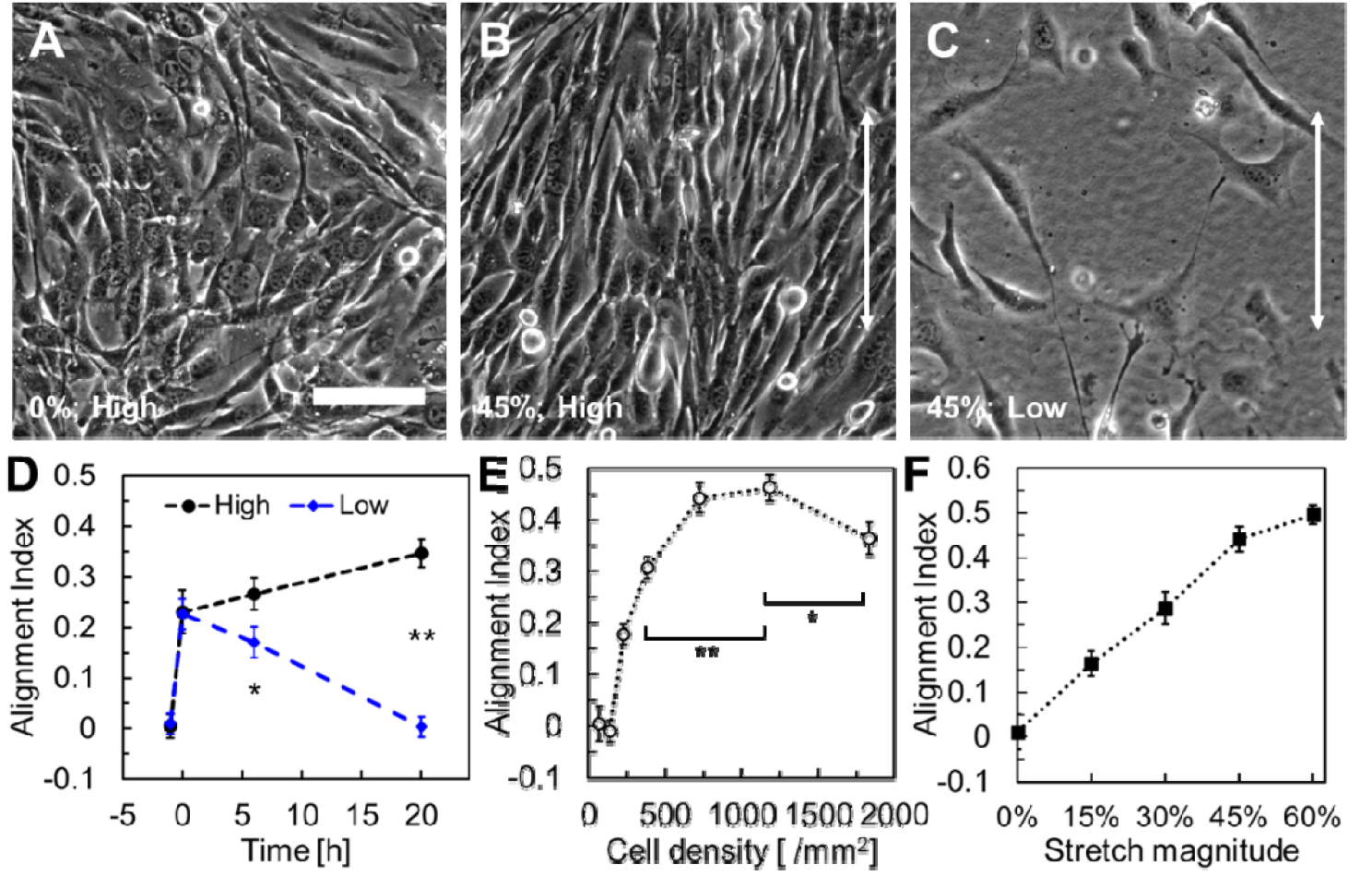
Mechanical stretch induced collective cell alignment. **A.** Random cell orientations at high cell density without mechanical stretch. **B**. Cell alignment with 45% static mechanical stretch at high cell density. **C**. Random cell orientations with 45% mechanical stretch at low cell density. **D**. Short-term and long-term cell alignment for high (570/mm^2^) and low (160/mm^2^) culture densities after a one-time 35% static stretch. *p=0.068; **p<10^-6^. n≥5. **E**. Effect of cell seeding density on cell alignment with 45% mechanical stretch. *p<0.05; **p<0.01. n≥3. **F**. Effect of stretch magnitude on cell alignment with a high cell density. In each condition, static stretches are applied three times evenly with a cell density of 730 cells/mm^2^ with different stretch magnitudes. n≥3. For each alignment index, analyzed cell number>1000. Scale bar: 100 μm.

We observe and quantify cell alignment 24 hours after the first stretch using the alignment index (see Methods). Initially near-confluent culture (725 cells/mm^2^) exhibits robust cell alignment (Fig. 1B and Fig. S1B), while sub-confluent culture (68 cells/mm^2^) does not (Fig. 1C and Fig. S1C), and cells do not align despite proliferating to a confluent density post-stretch (Fig. S1D). We quantify cell alignment immediately after stretch and again 20 hours later, revealing a *biphasic* alignment response: both high- and low-density cultures exhibit transient alignment shortly after stretching (Fig. 1D). However, over time, only high-density cultures maintain and further enhance alignment, while alignment in low-density cultures gradually declines. These results indicate that sustained alignment critically depends on pre-stretch cell density, with long-term alignment observed only under initially high-density conditions.

We next examine how loading conditions and cell density modulate long-term cell alignment (quantified at 24 hours of culture). Applying three cumulative 15% stretches with 45% total strain across all the cell densities, we find that long-term cell alignment increases with cell density until reaching a plateau, followed by a slight decline with further increasing cell density (Fig. 1E). To examine the effect of stretch magnitudes on cell alignment, high-density cultures are stretched with varying cumulative strains with evenly distributed stretch across three consecutive loadings in four-hour interval between each loading. We observe that cell alignment levels correlate strongly with stretch magnitude (Fig. 1F). These results demonstrate density-dependent collective cell alignment under static stretch: dense populations align robustly, while isolated cells do not.

### 3.2. Passive phase: substrate deformation drives transient cell alignment

To elucidate the mechanisms driving stretch-induced cell alignment, we first exclude potential factors. High-density cells failed to align on a pre-stretched substrate with the same stretch magnitude, regardless of whether fibronectin was coated before or after the stretch (Fig. S3). This rules out changes in substrate material properties, such as ECM fiber alignment[45-47], topography changes[48, 49], or stiffness anisotropy[17], as drivers of the observed alignment. In addition, soluble factors in the culture medium are also unlikely to be the reasons, as isotropic diffusion cannot generate large-scale anisotropic alignment patterns.

Studies have shown that substrate tension can direct immediate cell migration[50, 51] and increase the traction force[52]. Motivated by this, we asked whether C2C12 cells actively reorient toward the stretch direction after loading. To test this possibility, we quantify changes in cell orientation 2 and 4 h after a 15% stretch in low- and high-density cultures (Fig. S4). At low density, orientation changes remain centered near zero regardless of the initial cell orientations, indicating non-directional fluctuations rather than biased reorientation toward the stretch axis over this time scale. Thus, in this regime we do not attribute collective alignment to active, directionally biased mechanosensing at the single-cell level. In confluent cultures, the distribution deviates from symmetry, suggesting a degree of directional reorientation.

We next asked whether the short-term alignment can be explained by a purely passive mechanical mechanism. During stretch loading, cells deform together with the substrate, a phenomenon that we visualize by tracking fluorescent beads embedded beneath the monolayer (Fig. 2A-B). This *advective* substrate coupling (i.e., passive reorientation driven by the imposed substrate deformation rather than an active cellular response) biases cell orientation toward the stretch direction and can, in principle, induce alignment. To quantify this coupling, we measure both substrate deformation (from bead displacements) and cellular deformation (from bright-field cell boundaries) and defined a “deformation ratio” as the ratio of cellular to substrate deformation in the stretched direction (*y*), finding that cells exhibit nearly 1:1 strain matching with the substrate regardless of the initial cell orientations (Fig. 2C, black bars). Similarly, lateral compression along the *x* direction due to the Poisson effect also induces proportional shortening with a 1:1 deformation matching (Fig. 2C, blue bars), confirming bidirectional mechanical coupling.

**Figure 2.**
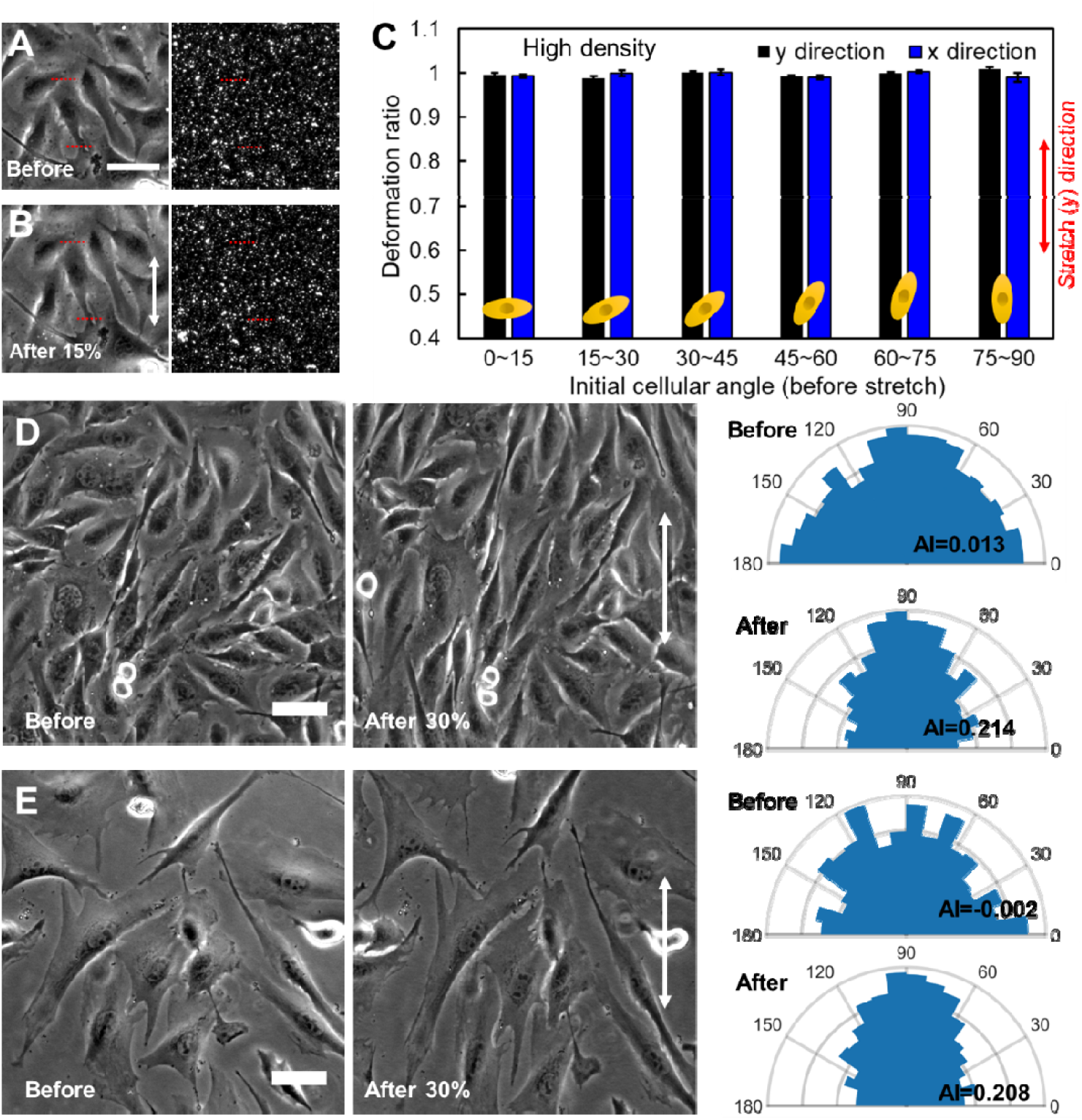
Deformation-initiated cell orientation bias and the resulting cell alignment. **A** and **B.** The position of a cell relative to the fluorescent beads-coated substrate before (A) and after (B) 15% uniaxial stretch. Red lines denote the top and bottom boundaries of the cell in the bright field image, corresponding to the locations in the fluorescent channel. **C**. Deformation ratio (defined in the Methods section) of cells in high density (590/mm^2^) after 15% mechanical stretch in terms of initial cell orientational angles in the y (stretched direction, red arrow) or x directions. n≥5 for each column. p>0.3 for each group compared with “1”. **D** and **E**. Immediate cell alignment as a result of 30% mechanical stretch in high (D, 620/mm^2^) and low (E, 130/mm^2^) cell densities. The right panels present rose plots for the cell angle distributions before and after the stretch, with the indicated alignment index. n>1000 for each rose-plot. Similar results were observed in n≥3 experiments. Scale bar: 50 μm.

To assess whether the measured substrate deformation is sufficient to reproduce the immediate post-stretch cell configuration, we computationally scale the pre-stretched cell image by the applied *y* and *x* strains to the substrate (Fig. S5A-B). The scaled image closely resembles the post-stretch bright field image of the cells immediately after stretch (Fig. S5C), indicating that cell shapes are passively advected with the substrate. The corresponding alignment indices before and immediately after stretch are quantified and suggest substrate deformation directly drives global cell alignment (Fig. 2D-E). These results demonstrate that short-term alignment arises from *passive* deformation: cells transiently mirror substrate strain without active reorganization.

### 3.3. Active phase: intercellular interactions drive long-term cell alignment

Substrate deformation-induced short-term alignment cannot explain the divergence in long-term alignment. In fact, substrate deformation is transmitted to cells similarly in both low and high-density cultures, as both show a deformation ratio close to unity (Fig. 2C and Fig. S6A). Additionally, the alignment response, reflected by cell orientational changes generated by stretch-induced deformation, depends on cell aspect ratio (Fig. S6B): cells with a larger aspect ratio are less responsive to orientation bias. We thus investigate whether cell aspect ratio depends on cell culture density. However, we find no significant difference in aspect ratios between low and high-density cultures (Fig. S6C). Since deformation ratio and aspect ratio, the key determinants of cell orientation bias under a given stretch, are similar across culture densities, the initial alignment levels immediately after a stretch are expected to be comparable, which is indeed the case (Fig. 2D-E and Fig. S6D). Therefore, the observed divergence in final alignment levels must arise from collective effects after the initial advective deformation. We thus propose that substrate deformation initiates transient cell alignment occurring for both dense and sparse cultures, which is then modified by density-dependent cooperative mechanisms, giving rise to the long-term alignment levels.

We next investigate the mechanisms underlying long-term cell alignment. Under 35% uniaxial stretch, cells in low-density cultures progressively lose their initial alignment induced by substrate deformation (Fig. 1D and Fig. S7A). In contrast, high-density cultures not only retain but further enhance their alignment over time (Fig. 1D and Fig. S7B). This density-dependent divergence highlights the importance of collective effects in sustaining alignment: while substrate deformation initiates cell alignment in both conditions, only dense cultures maintain and amplify it. Similar trends have been observed in collective durotaxis, where dense cell monolayers migrate directionally along shallow stiffness gradients, but isolated cells fail to respond. Such density-dependent sensitivity arises from cooperative cell-cell interactions, which are absent in sparse populations[26]. Our findings mirror this phenomenon, emphasizing that long-term alignment requires collective reinforcement, highlighting the critical role of intercellular cooperation in mediating sustained mechanical responses.

Cells in high-density cultures actively re-arrange their angles towards the stretched direction over time, progressively increasing their alignment. This behavior suggests a self-organization process that enhances alignment beyond the initial deformation-induced response. To dissect the underlying determinants, we examine cells initially polarized perpendicularly (*x*), which do not align passively to the stretch direction (*y*) via substrate deformation; note that this deformation alters their aspect ratios, but not their orientations. Notably, these cells can still reorient toward the stretch direction (*y*-axis) if neighboring cells are already aligned in that direction (Fig. 3A). This dependence on local context is consistent with prior observations of spontaneously formation of parallel arrays in confluent cell cultures[53, 54], where the presence of neighboring aligned cells leads to coherent nematic domains, resembling nematic grain boundaries (Fig. S8). In such cases, cells surrounded by misaligned neighbors, though adjectively aligned to some extent by the stretch, do not continue to reorient (Fig. 3B). These observations suggest that cell-cell contact within aligned regions actively promotes further alignment, as adopting a neighboring aligned configuration reduces the system’s free energy; re-orientation is unlikely to occur if none of the surrounding cells is aligned, as seen inside misaligned parallel arrays.

**Figure 3.**
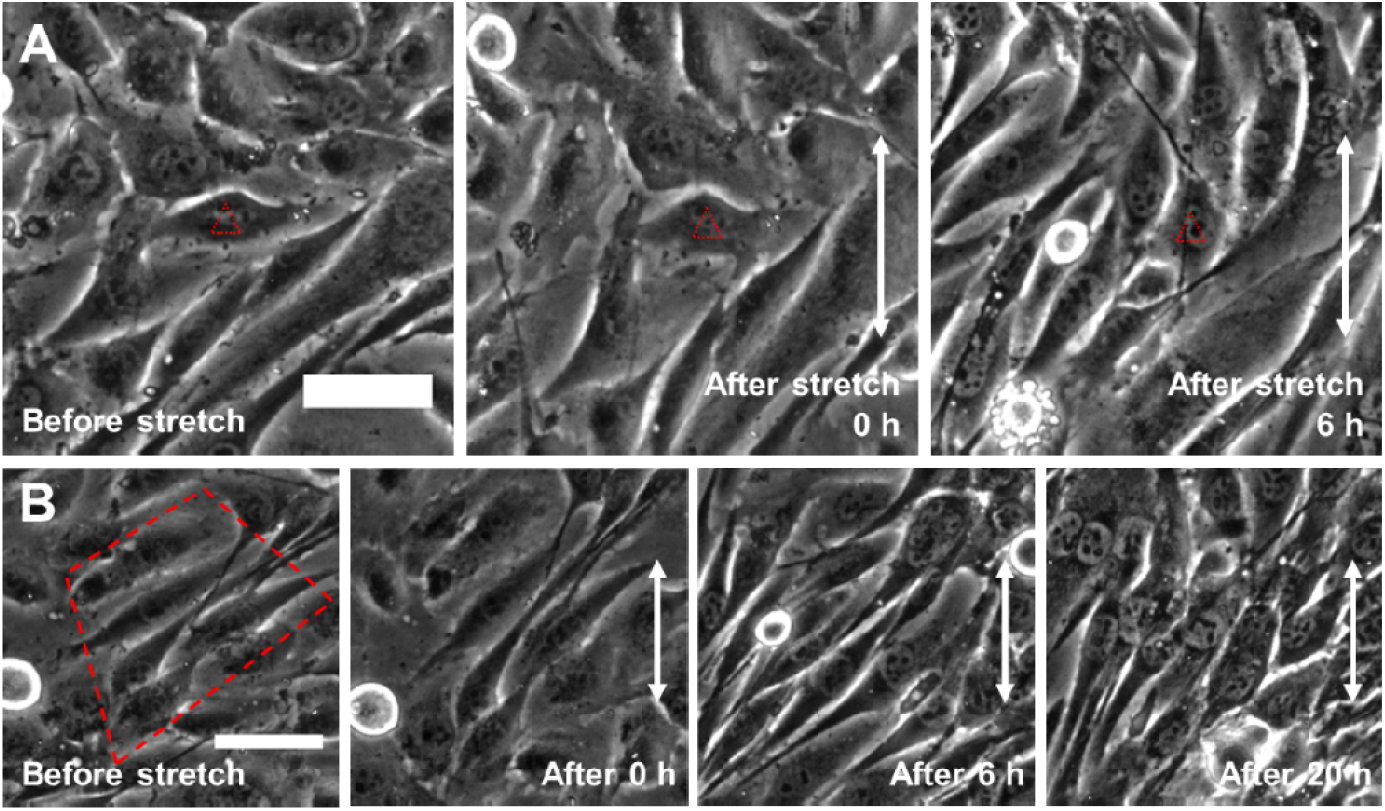
Collective features contributing to cell alignment. **A.** Cell angle change influenced by neighboring cells after a 30% stretch in high culture density (800/mm^2^). The representative cell, indicated by the red triangle, is initially oriented in the x direction thus could not passively reorient via substrate deformation. After 6 hours, this cell aligns in the stretched y direction (white arrow), a phenomenon that is accompanied by neighboring aligned cells. For n=12 cells that are x-direction oriented surrounded by partially aligned cells that we observe, all eventually show y-direction alignment. **B**. Representative image showing the inability to align for cells in a misaligned parallel arrays in high culture density (800/mm^2^). Among n=25 such parallel arrays analyzed, none show alignment, and all become nematic defects eventually. Scale bar: 50 μm.

To directly demonstrate the role of cell-cell interactions in cell alignment and to isolate this self-organization process from the initial stretch-induced alignment, we design a “secondary seeding” experiment (Fig. 4A). In this setup, we first seed cells at a near-confluent density (230/mm^2^) which are subjected to three cycles of stretching, totaling a 45% strain (Fig. 4B). This density is chose to minimize alignment loss over time, while giving space for secondary seeding. Immediately after the final stretch, a second group of cells, at the same density, is seeded into the same culture chamber, resulting in a high overall culture density. The second group of cells attaches to the substrate and spread within 1.5 hours, without overlapping on the initially seeded cells (Fig. 4C).

**Figure 4.**
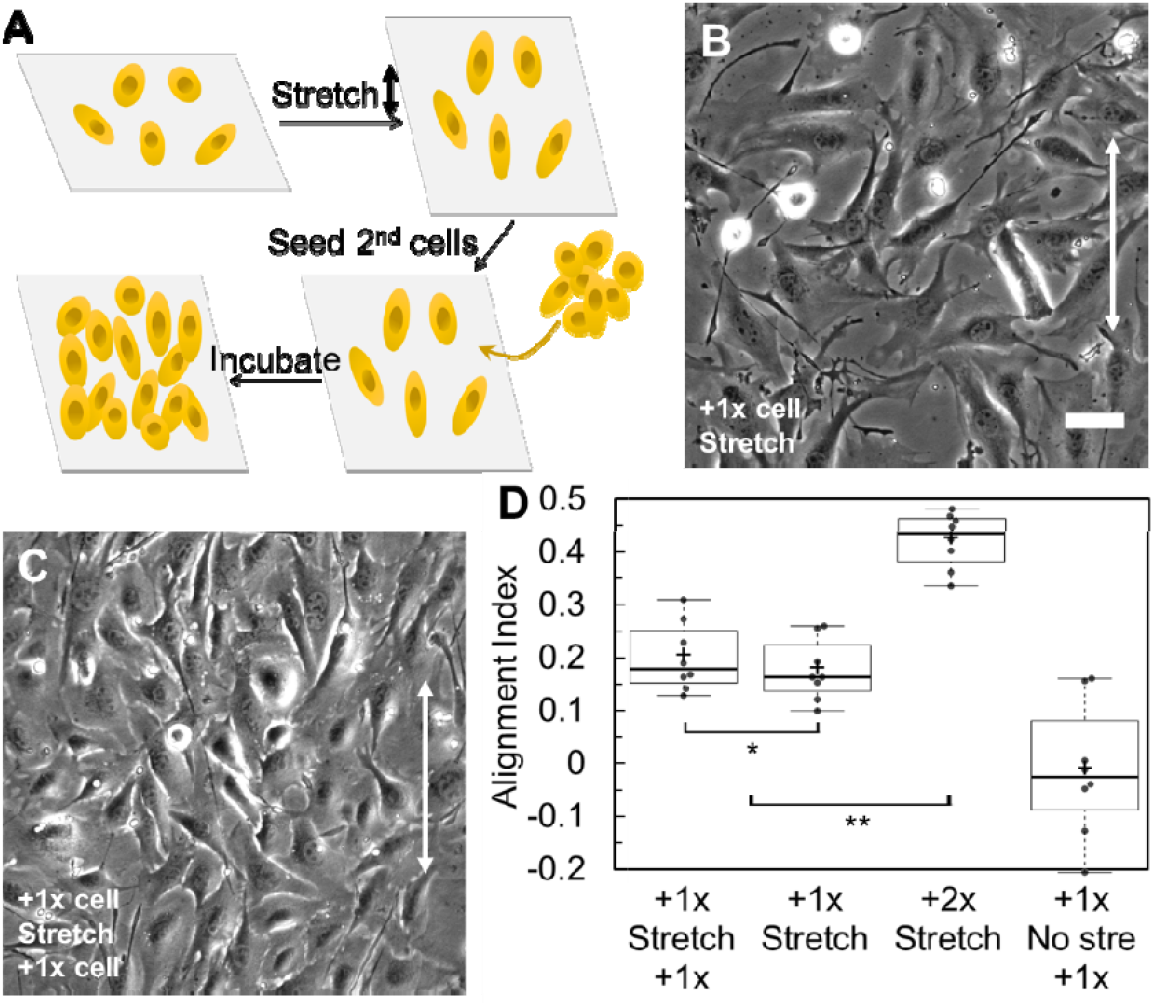
Self-organization process during cell alignment. **A.** Schematic of the “secondary seeding experiment” to distinguish the self-organization process from the initial alignment induced by uniaxial stretch. **B**. Representative bright field image of cells after 45% stretch. “+1x cell” indicates that cells are initially seeded at a near-confluent 1x density (230/mm^2^); “Stretch” indicates that the cells are subsequently subjected to a 45% stretch. **C**. Representative bright field image of the culture 1.5 hours after seeding the secondary cells. “+1x cell” below “Stretch” indicates that after stretching, an additional 1x density of cells is seeded into the culture. **D**. Quantification of cell alignment levels with or without secondary seeding after overnight incubation. First column: the “secondary seeding experiment” as described in C. Second column: no second group of cells is seeded after the stretch. Third column: initially 2x density of cells is seeded, and the stretch is applied, but no second group of cells is seeded. Last column: same as the “secondary seeding experiment” except that no stretch is applied. *p>0.1; **p<10^-5^. n≥5. Scale bar: 50 μm.

After overnight incubation, we observe that the overall alignment level of the entire cell population matches that of cultures seeded initially at near-confluent density without a secondary group (Fig. 4D and Fig. S9A-B). Although the alignment of the secondarily seeded cells could not be independently quantified, the comparable population-wide alignment suggests that the secondarily seeded cells align by following the orientation of the initially seeded cells, otherwise the final alignment level would have decreased by half. Note that cells seeded second could not align by residual substrate cues as pre-stretch does not cause cell alignment (Fig. S3). In contrast, when both groups are co-seeded simultaneously before stretching (i.e., one-time seeding at double the density), the final alignment is significantly higher, approximately twice that is observed in the secondary seeding experiment (Fig. 4D and Fig. S9C). This comparison demonstrates that the secondarily seeded cells aligned by following the orientation of their pre-aligned neighbors, as they have not directly experienced the stretch-induced alignment. We note that we do not differentially label the first and second batches of cells in the current study, and our analysis is therefore limited to population-averaged alignment. Future work using stable, non-transferable reporters to track the two populations separately will be valuable for resolving any subtle differences in their alignment dynamics. Together, our results confirm the existence of cell-cell interactions that enable cells to re-orient with their surrounding cells, resulting in self-organization behavior and the collective alignment.

### 3.4. Active alignment is kinetically driven by stochastic cellular forces

When cells are seeded onto a substrate, actomyosin motors generate intracellular contractile forces, which are transmitted both to the substrate, generating extracellular traction, and neighboring cells, generating intercellular tension. In isolated C2C12 cells under static stretch, traction force microscopy (TFM) shows that the overall traction magnitude and pattern change only modestly after loading (Fig. S10). This suggests that, in this system, the directionality and stochasticity of traction forces are not strongly modulated by static stretch. We note, however, that in other systems under dynamic (periodic) stretch, such as endothelial monolayers, traction field realignment can precede cell realignment and show pronounced fluidization-resolidification dynamics[55]. We next propose that the well-documented, orientation-randomizing “noise” generated by actomyosin pulses serves as a plausible effective source of stochasticity[56, 57]: in a coarse-grained sense, the variable directionality of cellular forces may act analogously to thermal fluctuations, enabling cells to probe their surroundings. An agent-based model is therefore used to examine the dynamic evolution of cell alignment. Ellipsoidal agents represent C2C12 cells and interact via a Gay-Berne potential that captures cell-cell adhesion and short-range repulsion[43], while a Langevin thermostat provides an effective stochastic forcing that mimics randomized active cellular forces (Fig. S11). Simulations are performed for four cases with identical parameters, differing only in initial orientation and cell density. In low-density cultures where cell-cell interactions are negligible, this stochastic forcing leads to gradual orientational randomization, yielding a disordered, orientation-entropy-dominated state (i.e., orientational disorder is favored, leading to a more isotropic distribution of cell angles) (Fig. 5A-B, Video S1). Conceptually, this effective-noise/activated reorientation picture is reminiscent of soft glassy rheology[58], where dynamics proceed via noise-assisted escape from metastable states in a disordered energy landscape. Similar soft-glassy phenomenology has also been invoked to interpret the scale-free rheology of adherent cells and the cytoskeleton[59]. In contrast, in high-density cultures where a cell is surrounded by nearly aligned neighbors, strong cell-cell interactions provide an energetic bias that stabilizes reorientation toward the local neighborhood. The resulting stable cell-cell junctions become less susceptible to stochastic fluctuations, thereby sustaining and enhancing alignment (Fig. 5C-D, Video S2).

**Figure 5.**
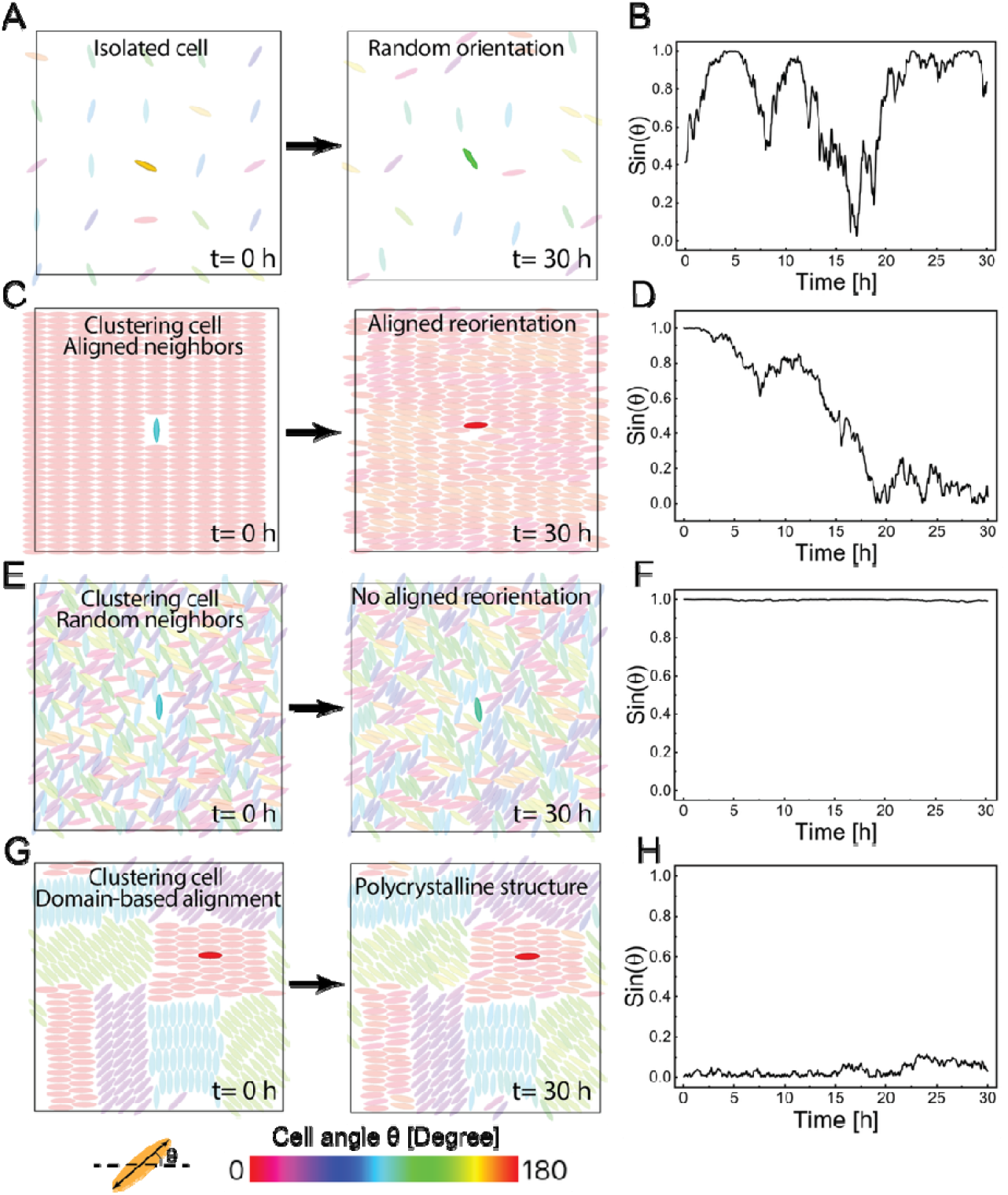
Agent-based simulations on cell alignment evolution driven by active cellular forces. **A.** Initial (left) and final (right) snapshot of low-density cultured cells. Entropy-dominant dynamics drive misalignment and the system shows random orientation. B. The single-cell orientation evolution reveals its random alignment. C. Initial (left) and final (right) snapshot of high-density cultured cells with aligned neighbors. The highlighted (green) cell is initially oriented at and does not mechanically interact with the surrounding cells at the initial time point. As interactions are established, cell-cell interactions act as an effective thermodynamic bias that tends to reinforce pre-introduced aligned configurations. D. The single-cell orientation evolution shows a realignment to the adjacent cells. **E**. Initial (left) and final (right) snapshot of high-density cultured cells with randomly aligned neighbors. In locally unaligned regions, cell reorientation is kinetically jammed by collective barriers and confinement, and reorientation is thermodynamically hindered due to non-aligned neighbors. **F**. The single-cell orientation evolution reveals a fixed misalignment because of local confinements. **G**. Initial (left) and final (right) snapshot of high-density cultured cells with a polycrystalline architecture: collective reorientation barriers kinetically trap cells in coherently misaligned domains, preventing global alignment. **H**. The single-cell orientation evolution shows a limited reorientation in a subdomain. The colormap shows the cell orientation angle, and the single-cell orientation evolution refers to the opaque cell of interest in the center of the simulation box. Each ellipsoid agent has a long axis of 100 μm.

The putative kinetic role of cellular forces in cell alignment is also evident from the incomplete alignment observed in high-density cultures experiments (Fig. S12). Under these conditions, cells experience resistance to reorientation from their tightly packed neighbors. This local deadlocking has been observed in agent-based modeling as well. In unaligned regions, dense local packing creates substantial reorientation barriers that kinetically trap cells in the unaligned orientations, as the self-generated cellular forces are insufficient to overcome the reorientation barriers imposed by surrounding cells (Fig. 5E-F, Video S3, and Fig. S12 red region). Note that the “energy wells/traps” in this schematic are meant in a coarse-grained, collective sense for multicellular orientational states, rather than as trapping of individual cytoskeletal constituents. Similarly, misaligned parallel arrays persist because reorienting entire nematic domains would require overcoming prohibitively high collective energy barriers (Fig. 3B). Perfect alignment would thus necessitate large-scale, coordinated domain reorientation, an event that appears kinetically unlikely at high densities (Fig. 5G-H, Video S4, and Fig. S12 green regions). As a result, the system resides in a metastable, higher-energy state without achieving global alignment. Together, we propose that alignment is dictated by the interplay between thermodynamic stabilization via intercellular interactions, active reorientation guided by neighbor cells, and kinetic resistance. This combined thermodynamic and agent-based framework explains the divergence of long-term alignment responses in dense and sparse cultures, as well as the polycrystalline-like domain structures observed in confluent tissues.

## 4. Discussion

While our current experiments identify the presence of intercellular interactions as a key ingredient for collective alignment, they do not allow us to discriminate between specific interacting mechanisms. Prior studies suggest that cells are able to communicate mechanically through elastic media[17, 60-62]. In our system, however, the available evidence argues against substrate-mediated cell-cell interactions as the dominant collective mechanism. For the PDMS membranes used in the static-stretch experiments, dividing the maximal cell traction stress (∼1 kPa) by the shear modulus of PDMS (∼1 MPa) yields a substrate strain of approximately 0.1%, far below the minimal threshold (∼2.5%) reported to be sensed by adjacent cells[61]. To further test whether cells communicate through traction forces on the soft substrate, we apply laser ablation experiments specifically designed to test whether substrate-mediated mechanical communication between neighboring cells plays a significant role in our system. We seed cells on a 2.6 kPa soft polyacrylamide hydrogel and incubate them with gold nanorods before the laser ablation experiments. To combine traction force microscopy (TFM) with laser ablation, we identify a pair of closely situated cells and capture an image of fluorescent beads underneath them (image I). Next, we open the laser shutter and apply a laser beam to ablate one of the cells for about 20 seconds. After laser ablation, we close the laser shutter and record the fluorescent beads underneath the cells (image II). Finally, we use a drop of sodium hydroxide solution to detach the cells and record the final image of the fluorescent beads (image III). By comparing images I and III, we obtain the traction force of the cell pair; by comparing images II and III, we obtain the traction force of the individual cell. Analysis of traction forces before and after ablation (Fig. S13) reveals no appreciable change in the traction pattern of the intact neighbor. Regarding time scales, we monitored the traction field over the minutes surrounding ablation, which is the typical window over which traction forces relax or reorganize after acute mechanical perturbations[55, 61, 63]. While we cannot rule out more subtle or longer-term effects, these results argue that force transmission via the soft substrate plays at most a minor role in mediating intercellular coupling in our experiments, and long-range substrate-mediated coupling is unlikely to be the dominant mechanism for the observed alignment.

These results motivate the exploration of alternative intercellular interaction mechanisms. Candidate mechanisms include cadherin-based cell-cell adhesion, contact guidance along pre-aligned cell bodies and aligned ECM deposited by the first layer, and purely geometric crowding/jamming in an anisotropic environment, any or all of which could drive the observed self-organization. For example, prior work has shown that weakening E-cadherin junctions can abrogate long-range orientational order in epithelia[64]. A critical next step will thus be to test whether analogous perturbations (e.g., N-cadherin knockdown or blocking antibodies), combined with time-resolved measurements of junctional maturation under stretch, alter the density-dependent alignment reported here. Likewise, purely physical crowding may contribute: high packing can suppress orientational randomization and promote local co-alignment through steric/geometric constraints, thereby reinforcing the stretch-imprinted bias even in the absence of strong junctional mechanotransductive signaling. At the same time, crowding can induce jammed dynamics that slow neighbor rearrangements and defect coarsening, allowing misaligned domains to persist as a polycrystalline-like, metastable architecture[65]. Such crowding/jamming effects could arise even in the absence of strong mechanotransductive or cadherin-based signaling[66, 67]. Together, these considerations highlight that future work should explicitly dissect the molecular and physical identity of the intercellular coupling. For this reason, we present the thermodynamic and kinetic picture (Fig. 5) as a conceptual framework consistent with our current data, and systematic perturbations of cell-cell interactions will be essential to test and refine this framework.

In our study, transient mechanical stretch serves as a time-limited stimulus to probe cellular memory formation through dissipative adaptation, a nonequilibrium process where energy dissipation drives functional reorganization. Low-density cultures exhibit short-term alignment in response to stretching (passive phase), but gradually revert to disordered states post-stimulus, reflecting a lack of lasting mechanical memory. In contrast, high-density cultures retain long-term alignment, demonstrating collective mechanical memory. This memory emerges via a dissipative adaptation cycle: cellular forces do work to dissipate energy, which is fueled by actomyosin activity and reinforces cell-cell adhesion; the reinforced adhesion network “records” the stretch direction, creating a metastable configuration resistant to noise, i.e., fluctuating cellular forces. This process is dissipative: energy consumed by cellular forces and intercellular interactions remodeling is irreversibly lost, yet it enables adaptation by “locking” the system into the aligned state. Thus, mechanical memory in tissues operates as a dissipative adaptation phenomenon, where irreversible processes convert transient stimuli into persistent structural order.

In this study, we focused specifically on C2C12 myoblast monolayers subjected to static (stepwise) uniaxial stretch, and our conclusions about biphasic, density-dependent alignment are therefore demonstrated within this system. Static or quasi-static uniaxial loading has similarly been shown to align other mesoderm-derived, spindle-shaped cells, such as C2C12 myotubes and fibroblast-populated collagen lattices, parallel to the loading axis, suggesting that related mechanisms may operate in contractile mesenchymal cell types that share elongated morphology and strong traction generation [8, 31, 32]. By contrast, many epithelial, endothelial, and fibroblast systems subjected to cyclic uniaxial stretch reorient nearly perpendicular to the stretch direction in a manner that depends sensitively on strain amplitude and frequency, indicating that the loading waveform can fundamentally reshape the alignment outcome[68, 69]. We therefore anticipate that the balance between the passive, deformation-driven phase and the active, interaction-mediated phase will depend not only on cell lineage and morphology (e.g., mesodermal myoblasts and fibroblasts versus epithelial/endothelial cells) but also on the detailed stretch protocol (static versus cyclic, magnitude, and history). Systematic comparative studies that vary cell type and mechanical regimen will be required to test how broadly the biphasic, density-dependent framework identified here extends beyond C2C12 myoblasts. In addition, large transient deformations have been reported to fluidize and reconfigure cytoskeletal networks [70], suggesting that stretch waveform/pulsatility could, in principle, facilitate escape from kinetically trapped misaligned domains; testing pulse-release protocols is an interesting direction for future work.

## 5. Conclusions

In this study, we demonstrate a collective mode of cell alignment, where high-density C2C12 cultures align robustly, while low-density cultures do not. The alignment process proceeds in a biphasic manner: a short-term passive phase, where substrate deformation transiently biases cell orientations along the stretch direction, and a long-term active phase governed by cell-cell interactions. Notably, both dense and sparse cultures exhibit similar transient alignment after stretch, allowing us to isolate the mechanisms responsible for the long-term, density-dependent divergence that follows. Overtime, only high-density cultures maintain and enhance alignment, and this sustained alignment arises from two key features: (1) strong cell-cell adhesions thermodynamically stabilize aligned states, and (2) cells reorient toward the stretch direction when surrounded by already-aligned neighbors. In contrast, sparse cultures lack sufficient adhesion and neighbor-induced guidance, allowing stochastic cellular forces to disrupt alignment over time. Our findings provide a combined thermodynamic and kinetic framework in collective cell alignment, and at the same time suggest a simple, scalable strategy for aligning cells in tissue engineering, particularly for skeletal muscle regeneration.

## CRediT authorship contribution statement

**Xuechen Shi**: Conceptualization, Methodology, Investigation, Formal analysis, Visualization, Writing - original draft, Writing - review & editing

**Luyi Feng**: Methodology, Investigation, Formal analysis, Visualization, Writing - review & editing

**Sulin Zhang**: Conceptualization, Resources, Funding acquisition, Project administration, Supervision, Writing - original draft, Writing - review & editing

## Acknowledgements

This research was supported by the National Institutes of Health (NHLBI R21 HL122902) and National Science Foundation (NSF: CMMI-0754463/0644599, CMMI 1933398).

## References

[1] M. Wakelam, The fusion of myoblasts, Biochemical Journal 228(1) (1985) 1.

[2] M. Junkin, S.L. Leung, S. Whitman, C.C. Gregorio, P.K. Wong, Cellular self-organization by autocatalytic alignment feedback, Journal of cell science 124(24) (2011) 4213–4220.

[3] A.-C. Vion, M. Kheloufi, A. Hammoutene, J. Poisson, J. Lasselin, C. Devue, I. Pic, N. Dupont, J. Busse, K. Stark, Autophagy is required for endothelial cell alignment and atheroprotection under physiological blood flow, Proceedings of the National Academy of Sciences 114(41) (2017) E8675–E8684.

[4] D.E. Conway, M.A. Schwartz, Flow-dependent cellular mechanotransduction in atherosclerosis, Journal of cell science 126(22) (2013) 5101–5109.

[5] S. Vyalov, B. Langille, A.I. Gotlieb, Decreased blood flow rate disrupts endothelial repair in vivo, The American journal of pathology 149(6) (1996) 2107.

[6] J.H. Wang, F. Jia, T.W. Gilbert, S.L. Woo, Cell orientation determines the alignment of cell-produced collagenous matrix, Journal of biomechanics 36(1) (2003) 97–102.

[7] Z. Yin, X. Chen, J.L. Chen, W.L. Shen, T.M.H. Nguyen, L. Gao, H.W. Ouyang, The regulation of tendon stem cell differentiation by the alignment of nanofibers, Biomaterials 31(8) (2010) 2163–2175.

[8] K. Chen, A. Vigliotti, M. Bacca, R.M. McMeeking, V.S. Deshpande, J.W. Holmes, Role of boundary conditions in determining cell alignment in response to stretch, Proceedings of the National Academy of Sciences 115(5) (2018) 986–991.

[9] S. Kalinina, H. Gliemann, M. López-García, A. Petershans, J. Auernheimer, T. Schimmel, M. Bruns, A. Schambony, H. Kessler, D. Wedlich, Isothiocyanate-functionalized RGD peptides for tailoring cell-adhesive surface patterns, Biomaterials 29(20) (2008) 3004–3013.

[10] C. Williams, Y. Tsuda, B.C. Isenberg, M. Yamato, T. Shimizu, T. Okano, J.Y. Wong, Aligned cell sheets grown on thermo-responsive substrates with microcontact printed protein patterns, Adv Mater 21(21) (2009) 2161–2164.

[11] J.S. Choi, S.J. Lee, G.J. Christ, A. Atala, J.J. Yoo, The influence of electrospun aligned poly (epsilon-caprolactone)/collagen nanofiber meshes on the formation of self-aligned skeletal muscle myotubes, Biomaterials 29(19) (2008) 2899–2906.

[12] J.I. Andersen, C.P. Pennisi, T. Fink, V. Zachar, Focal adhesion kinase activation is necessary for stretch-induced alignment and enhanced differentiation of myogenic precursor cells, Tissue Engineering Part A 24(7-8) (2018) 631–640.

[13] K. Kurpinski, J. Chu, C. Hashi, S. Li, Anisotropic mechanosensing by mesenchymal stem cells, Proceedings of the National Academy of Sciences 103(44) (2006) 16095–16100.

[14] A. Livne, E. Bouchbinder, B. Geiger, Cell reorientation under cyclic stretching, Biophysical journal 106(2) (2014) 42a.

[15] R. Kaunas, P. Nguyen, S. Usami, S. Chien, Cooperative effects of Rho and mechanical stretch on stress fiber organization, Proceedings of the National Academy of Sciences 102(44) (2005) 15895–15900.

[16] Y. Li, G. Huang, X. Zhang, L. Wang, Y. Du, T.J. Lu, F. Xu, Engineering cell alignment in vitro, Biotechnology advances 32(2) (2014) 347–365.

[17] I.B. Bischofs, U.S. Schwarz, Cell organization in soft media due to active mechanosensing, Proceedings of the National Academy of Sciences 100(16) (2003) 9274–9279.

[18] R. De, A. Zemel, S.A. Safran, Dynamics of cell orientation, Nature Physics 3(9) (2007) 655–659.

[19] A. Ray, O. Lee, Z. Win, R.M. Edwards, P.W. Alford, D.-H. Kim, P.P. Provenzano, Anisotropic forces from spatially constrained focal adhesions mediate contact guidance directed cell migration, Nature communications 8(1) (2017) 14923.

[20] J.H.-C. Wang, P. Goldschmidt-Clermont, J. Wille, F.C.-P. Yin, Specificity of endothelial cell reorientation in response to cyclic mechanical stretching, Journal of biomechanics 34(12) (2001) 1563–1572.

[21] D. Ellison, A. Mugler, M.D. Brennan, S.H. Lee, R.J. Huebner, E.R. Shamir, L.A. Woo, J. Kim, P. Amar, I. Nemenman, Cell-cell communication enhances the capacity of cell ensembles to sense shallow gradients during morphogenesis, Proceedings of the National Academy of Sciences 113(6) (2016) E679–E688.

[22] S. Fancher, A. Mugler, Fundamental limits to collective concentration sensing in cell populations, Physical review letters 118(7) (2017) 078101.

[23] A. Mugler, A. Levchenko, I. Nemenman, Limits to the precision of gradient sensing with spatial communication and temporal integration, Proceedings of the National Academy of Sciences 113(6) (2016) E689–E695.

[24] G. Malet-Engra, W. Yu, A. Oldani, J. Rey-Barroso, N.S. Gov, G. Scita, L. Dupré, Collective cell motility promotes chemotactic prowess and resistance to chemorepulsion, Current Biology 25(2) (2015) 242–250.

[25] E. Theveneau, L. Marchant, S. Kuriyama, M. Gull, B. Moepps, M. Parsons, R. Mayor, Collective chemotaxis requires contact-dependent cell polarity, Dev Cell 19(1) (2010) 39–53.

[26] R. Sunyer, V. Conte, J. Escribano, A. Elosegui-Artola, A. Labernadie, L. Valon, D. Navajas, J.M. García-Aznar, J.J. Muñoz, P. Roca-Cusachs, Collective cell durotaxis emerges from long-range intercellular force transmission, Science 353(6304) (2016) 1157–1161.

[27] N.D. Bade, R.D. Kamien, R.K. Assoian, K.J. Stebe, Curvature and Rho activation differentially control the alignment of cells and stress fibers, Science advances 3(9) (2017) e1700150.

[28] M.S. Grigola, C.L. Dyck, D.S. Babacan, D.N. Joaquin, K.J. Hsia, Myoblast alignment on 2D wavy patterns: Dependence on feature characteristics and cell - cell interaction, Biotechnology and bioengineering 111(8) (2014) 1617–1626.

[29] S. Coyle, B. Doss, Y. Huo, H.R. Singh, D. Quinn, K.J. Hsia, P.R. LeDuc, Cell alignment modulated by surface nano-topography-Roles of cell-matrix and cell-cell interactions, Acta Biomaterialia 142 (2022) 149–159.

[30] Y. Zhang, X. Shi, T. Zhao, C. Huang, Q. Wei, X. Tang, L.C. Santy, M.T.A. Saif, S. Zhang, A traction force threshold signifies metastatic phenotypic change in multicellular epithelia, Soft Matter 15(36) (2019) 7203–7210.

[31] A.M. Collinsworth, C.E. Torgan, S.N. Nagda, R.J. Rajalingam, W.E. Kraus, G.A. Truskey, Orientation and length of mammalian skeletal myocytes in response to a unidirectional stretch, Cell and tissue research 302(2) (2000) 243–251.

[32] M. Eastwood, V. Mudera, D. McGrouther, R. Brown, Effect of precise mechanical loading on fibroblast populated collagen lattices: morphological changes, Cell motility and the cytoskeleton 40(1) (1998) 13–21.

[33] N. Zhu, H.K. Kwong, Y. Bao, T.-H. Chen, Chiral orientation of skeletal muscle cells requires rigid substrate, Micromachines 8(6) (2017) 181.

[34] Y. Huang, Y. Bao, H.K. Kwong, T.H. Chen, M.L. Lam, Outline-etching image segmentation reveals enhanced cell chirality through intercellular alignment, Biotechnology and Bioengineering 115(10) (2018) 2595–2603.

[35] Y. Pang, X. Wang, D. Lee, H.P. Greisler, Dynamic quantitative visualization of single cell alignment and migration and matrix remodeling in 3-D collagen hydrogels under mechanical force, Biomaterials 32(15) (2011) 3776–3783.

[36] F. Xu, T. Beyazoglu, E. Hefner, U.A. Gurkan, U. Demirci, Automated and adaptable quantification of cellular alignment from microscopic images for tissue engineering applications, Tissue Engineering Part C: Methods 17(6) (2011) 641–649.

[37] M. Sun, A.B. Bloom, M.H. Zaman, Rapid quantification of 3D collagen fiber alignment and fiber intersection correlations with high sensitivity, PLoS One 10(7) (2015) e0131814.

[38] S.R.K. Vedula, G. Peyret, I. Cheddadi, T. Chen, A. Brugués, H. Hirata, H. Lopez-Menendez, Y. Toyama, L. Neves de Almeida, X. Trepat, Mechanics of epithelial closure over non-adherent environments, Nature communications 6(1) (2015) 6111.

[39] L. Casares, R. Vincent, D. Zalvidea, N. Campillo, D. Navajas, M. Arroyo, X. Trepat, Hydraulic fracture during epithelial stretching, Nat Mater 14(3) (2015) 343–51.

[40] Q. Wei, X. Shi, T. Zhao, P. Cai, T. Chen, Y. Zhang, C. Huang, J. Yang, X. Chen, S. Zhang, Actin-ring segment switching drives nonadhesive gap closure, Proc Natl Acad Sci U S A 117(52) (2020) 33263–33271.

[41] X.C. Shi, Z.Z. Liu, L.Y. Feng, T.K. Zhao, C.Y. Hui, S.L. Zhang, Elastocapillarity at Cell-Matrix Contacts, Phys Rev X 12(2) (2022).

[42] S. Plimpton, Fast parallel algorithms for short-range molecular dynamics, Journal of computational physics 117(1) (1995) 1–19.

[43] W.M. Brown, M.K. Petersen, S.J. Plimpton, G.S. Grest, Liquid crystal nanodroplets in solution, The Journal of chemical physics 130(4) (2009).

[44] B. Dünweg, W. Paul, Brownian dynamics simulations without Gaussian random numbers, International Journal of Modern Physics C 2(03) (1991) 817–827.

[45] H.H. Vandenburgh, P. Karlisch, Longitudinal growth of skeletal myotubes in vitro in a new horizontal mechanical cell stimulator, In vitro cellular & developmental biology 25(7) (1989) 607–616.

[46] H.H. Vandenburgh, Dynamic mechanical orientation of skeletal myofibers in vitro, Developmental Biology 93(2) (1982) 438–443.

[47] A. Matsugaki, N. Fujiwara, T. Nakano, Continuous cyclic stretch induces osteoblast alignment and formation of anisotropic collagen fiber matrix, Acta Biomaterialia 9(7) (2013) 7227–7235.

[48] D.J. Evans, S. Britland, P.M. Wigmore, Differential response of fetal and neonatal myoblasts to topographical guidance cues in vitro, Development genes and evolution 209 (1999) 438–442.

[49] P.Y. Wang, H.T. Yu, W.B. Tsai, Modulation of alignment and differentiation of skeletal myoblasts by submicron ridges/grooves surface structure, Biotechnology and bioengineering 106(2) (2010) 285–294.

[50] C.-M. Lo, H.-B. Wang, M. Dembo, Y.-l. Wang, Cell movement is guided by the rigidity of the substrate, Biophysical journal 79(1) (2000) 144–152.

[51] H.-B. Wang, M. Dembo, S.K. Hanks, Y.-l. Wang, Focal adhesion kinase is involved in mechanosensing during fibroblast migration, Proceedings of the National Academy of Sciences 98(20) (2001) 11295–11300.

[52] S. Munevar, Y.-l. Wang, M. Dembo, Regulation of mechanical interactions between fibroblasts and the substratum by stretch-activated Ca2+ entry, Journal of cell science 117(1) (2004) 85–92.

[53] L. Edelstein-Keshet, G.B. Ermentrout, Models for contact-mediated pattern formation: cells that form parallel arrays, Journal of mathematical biology 29 (1990) 33–58.

[54] L. Petitjean, M. Reffay, E. Grasland-Mongrain, M. Poujade, B. Ladoux, A. Buguin, P. Silberzan, Velocity fields in a collectively migrating epithelium, Biophysical journal 98(9) (2010) 1790–1800.

[55] R. Krishnan, E.P. Canović, A.L. Iordan, K. Rajendran, G. Manomohan, A.P. Pirentis, M.L. Smith, J.P. Butler, J.J. Fredberg, D. Stamenović, Fluidization, resolidification, and reorientation of the endothelial cell in response to slow tidal stretches, American Journal of Physiology-Cell Physiology 303(4) (2012) C368–C375.

[56] A.R. Harris, L. Peter, J. Bellis, B. Baum, A.J. Kabla, G.T. Charras, Characterizing the mechanics of cultured cell monolayers, Proceedings of the National Academy of Sciences 109(41) (2012) 16449–16454.

[57] N. Khalilgharibi, J. Fouchard, N. Asadipour, R. Barrientos, M. Duda, A. Bonfanti, A. Yonis, A. Harris, P. Mosaffa, Y. Fujita, Stress relaxation in epithelial monolayers is controlled by the actomyosin cortex, Nature physics 15(8) (2019) 839–847.

[58] P. Sollich, F. Lequeux, P. Hébraud, M.E. Cates, Rheology of soft glassy materials, Physical review letters 78(10) (1997) 2020.

[59] B. Fabry, G.N. Maksym, J.P. Butler, M. Glogauer, D. Navajas, J.J. Fredberg, Scaling the microrheology of living cells, Physical review letters 87(14) (2001) 148102.

[60] J.P. Winer, S. Oake, P.A. Janmey, Non-linear elasticity of extracellular matrices enables contractile cells to communicate local position and orientation, PloS one 4(7) (2009) e6382.

[61] X. Tang, P. Bajaj, R. Bashir, T.A. Saif, How far cardiac cells can see each other mechanically, Soft Matter 7(13) (2011) 6151–6158.

[62] C.A. Reinhart-King, M. Dembo, D.A. Hammer, Cell-cell mechanical communication through compliant substrates, Biophysical journal 95(12) (2008) 6044–6051.

[63] R. Krishnan, C.Y. Park, Y.-C. Lin, J. Mead, R.T. Jaspers, X. Trepat, G. Lenormand, D. Tambe, A.V. Smolensky, A.H. Knoll, Reinforcement versus fluidization in cytoskeletal mechanoresponsiveness, PloS one 4(5) (2009) e5486.

[64] B. Ladoux, R.M. Mege, Mechanobiology of collective cell behaviours, Nat Rev Mol Cell Biol 18(12) (2017) 743–757.

[65] L. Atia, J.J. Fredberg, N.S. Gov, A.F. Pegoraro, Are cell jamming and unjamming essential in tissue development?, Cells & development 168 (2021) 203727.

[66] M. Sadati, N.T. Qazvini, R. Krishnan, C.Y. Park, J.J. Fredberg, Collective migration and cell jamming, Differentiation 86(3) (2013) 121–125.

[67] E. Lawson-Keister, M.L. Manning, Jamming and arrest of cell motion in biological tissues, Current Opinion in Cell Biology 72 (2021) 146–155.

[68] J.-C. Lien, Y.-l. Wang, Cyclic stretching-induced epithelial cell reorientation is driven by microtubule-modulated transverse extension during the relaxation phase, Scientific Reports 11(1) (2021) 14803.

[69] U. Faust, N. Hampe, W. Rubner, N. Kirchgessner, S. Safran, B. Hoffmann, R. Merkel, Cyclic stress at mHz frequencies aligns fibroblasts in direction of zero strain, PloS one 6(12) (2011) e28963.

[70] P. Bursac, G. Lenormand, B. Fabry, M. Oliver, D.A. Weitz, V. Viasnoff, J.P. Butler, J.J. Fredberg, Cytoskeletal remodelling and slow dynamics in the living cell, Nature materials 4(7) (2005) 557–561.

