## Supplementary Figures for "Static mechanical stretch induces collective alignment of C2C12 myoblasts"

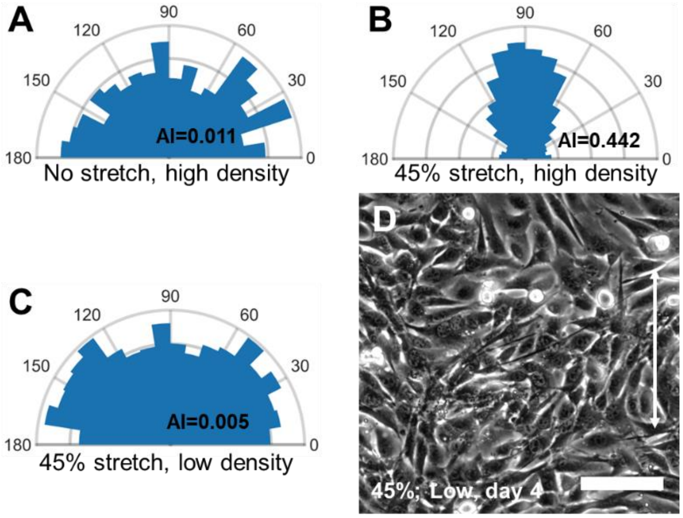


**Figure S1**. Collective cell alignment induced by mechanical stretch. **A**. Rose plot of cell orientation angles without static mechanical stretch in high culture density. **B**. Rose plot of cell orientation angles with 45% static mechanical stretch in high culture density. **C**. Rose plot of cell orientation angles with 45% static mechanical stretch in low culture density. **D**. Random cell orientations with 45% mechanical stretch at low cell density, taken three days after stretch once cells become fully confluent. n>1000 for each rose plot. Scale bar: 100 µm.

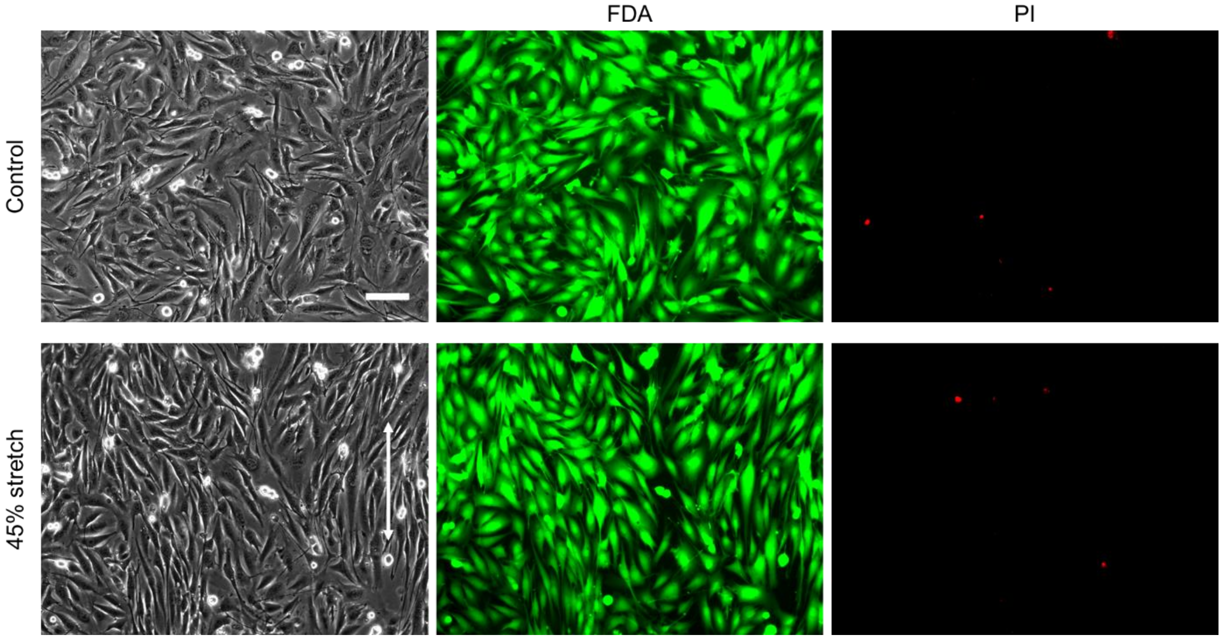


**Figure S2**. Impact of mechanical stretch on cell viability. Cells after stretch are stained with 8 µg/ml fluorescein diacetate (FDA) and 20 µg/ml propidium iodide (PI) for 5 minutes. Cells maintain viability after three stretches with a total magnitude of 45%. Scale bar: 100 µm.

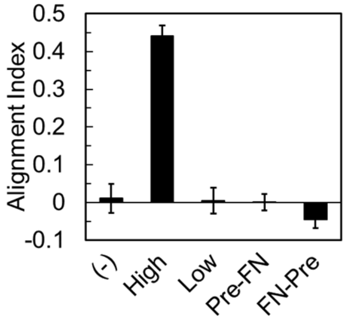


**Figure S3**. Average alignment indexes under different conditions. (-) indicates no stretch at high cell density. High: high cell density with 45% stretch. $p<0.001$ between "High" and all other groups. Low: low cell density with 45% stretch. Pre-FN: pre-stretch of the substrate before fibronectin coating prior to cell seeding at high density. FN-Pre: pre-stretch of the substrate after fibronectin coating prior to cell seeding at high density. $n \geq 3$ for each condition. For each alignment index, the analyzed cell number>1000.

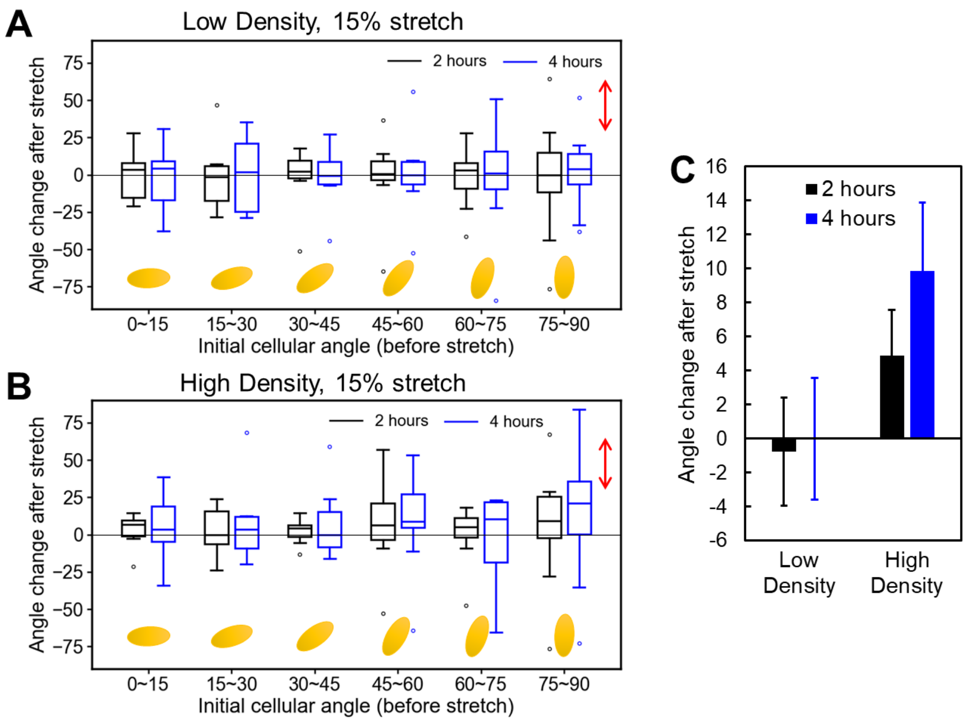


**Figure S4**. Post-stretch orientation changes in low- and high-density cultures after 2 or 4 hours of 15% mechanical stretch. Here, the cell orientation angle $\theta$ (-90° to 90°) was defined as the angle between the major axis of the cell body and the x-axis (perpendicular to the stretch axis); note that this definition differs from the cell angle (0° to 180°) used elsewhere. With this convention, a larger absolute angle $|\theta|$ corresponds to stronger alignment with the stretch axis. We therefore used the angle change $\Delta|\theta|$ to quantify reorientation: positive angle change $\Delta|\theta| > 0$ indicates rotation toward the stretch axis, whereas $\Delta|\theta| < 0$ indicates rotation away. Further, to examine the effects of initial cell angle, cells were grouped by their pre-stretch orientation. **A**. Angle change in low cell density at 70/mm$^2$. $\Delta|\theta|$ is symmetric about zero across all bins, indicating no directional drift. **B**. Active angle change in high cell density at 650/mm$^2$. n≥10 for each bin. Red arrows indicate the stretch direction (y), and yellow ellipses indicate the initial cell orientations. **C**. Average angle change for both high and low cell densities across all analyzed cells. n>80 for each bar. No statistically significant differences are observed (p>0.18 for 2 h and p=0.07 for 4 h between Low and High Density groups).

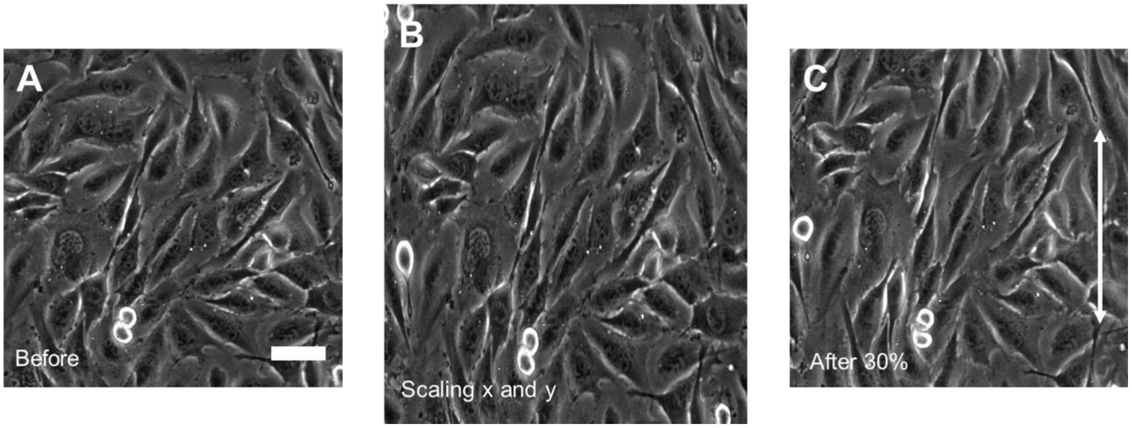


**Figure S5**. Image scaling analysis illustrating deformation-induced passive mode in the short-term response. **A**. Original bright field image of cells. **B**. Computationally scaled image from A by -10% in the x direction and +30% in the y direction. **C**. Bright field image of the same region immediately after a 30% mechanical stretch. The close visual similarity between B and C indicates that cells transiently mirror the substrate strain after stretch. Scale bar: 50 µm.

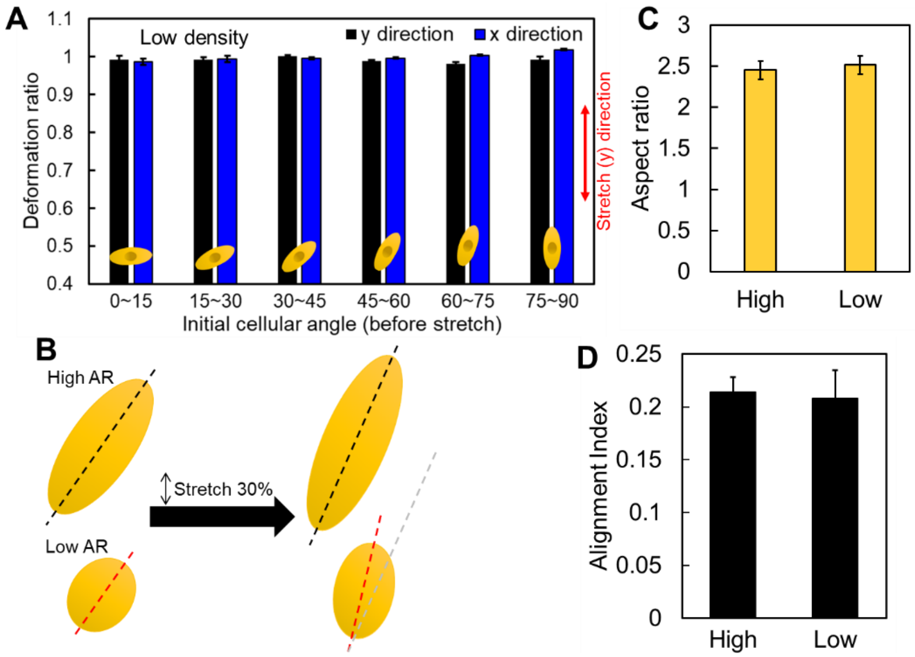


**Figure S6**. Deformation-induced cell alignment in high and low densities. **A**. Deformation ratio of cells in low density after 15% mechanical stretch in terms of initial cell orientation angles in the y (stretched direction, red arrow) or x directions. n≥5 for each column. **B**. Dependence of deformation-induced cell alignment on aspect ratio (AR). Cells with low aspect ratios are more prone to align given the same deformation. **C**. Cell aspect ratios in high (650/mm$^2$) and low (70/mm$^2$) cell densities. n>90 for each column. No statistically significant difference is found (p=0.69). **D**. Cell alignment levels immediately after 30% uniaxial stretch in high (620/mm$^2$) and low (130/mm$^2$) cell densities. No statistically significant difference is found (p=0.86). n≥4.

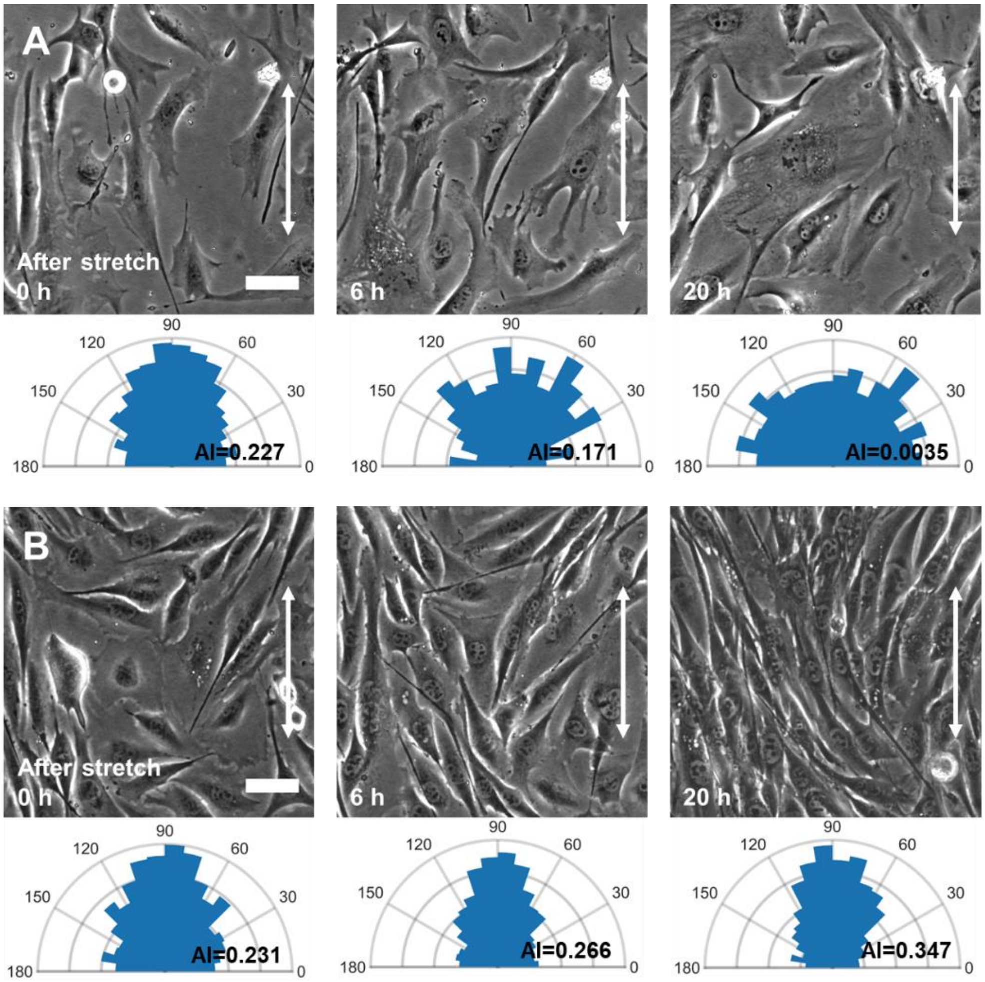


**Figure S7**. Cell alignment levels over time revealing the collective features during alignment. **A**. Loss of alignment in cells with low culture density (160/mm$^2$) over time. **B**. Enhancement of alignment in cells with high culture density (570/mm$^2$) over time. A&B: Stretch magnitude is 35%. Each rose plot underneath the image represents the cell angle distribution, with the alignment index indicated. n>1000 for each plot. Scale bar: 50 μm.

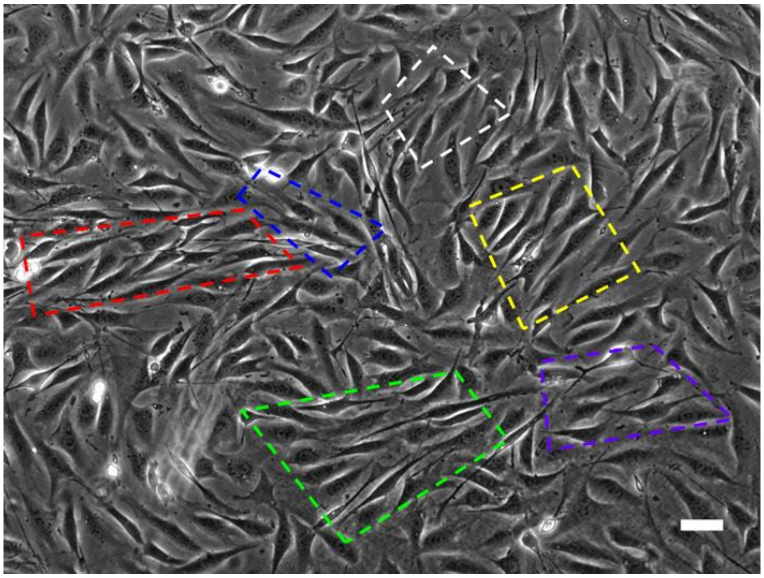

**Figure S8**. Multiple oriented domains in a cell culture landscape. Each oriented domain is indicated with a dashed quadrangle. Scale bar: 50 μm.

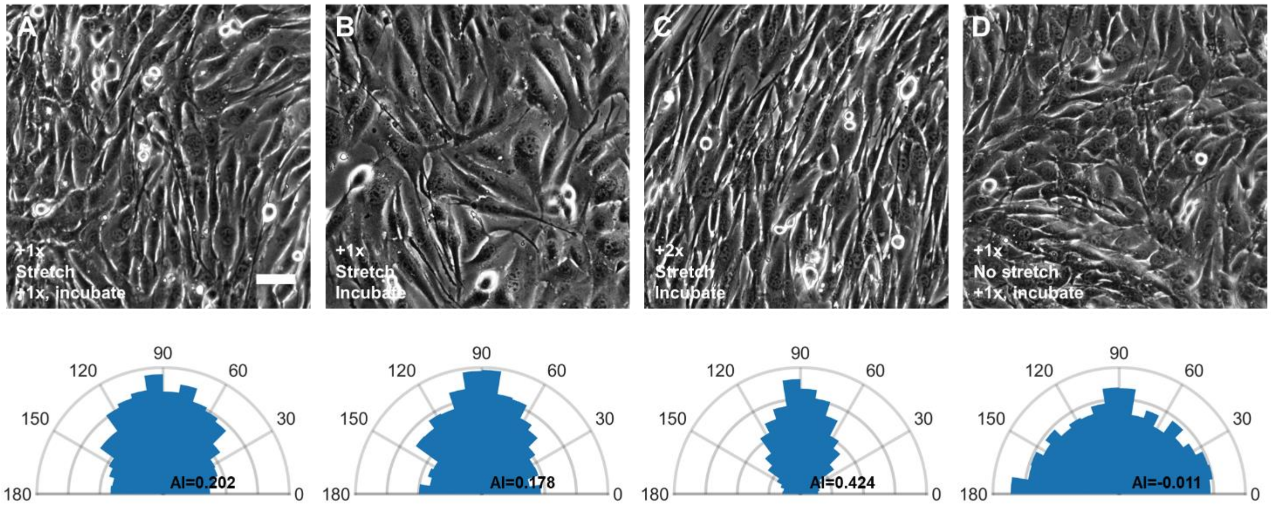


**Figure S9**. Representative images for the "secondary seeding experiment". The experimental condition for each image is explained in Figure 4. Images are taken after overnight incubation. Each rose plot underneath the image represents the cell angle distribution, with alignment index indicated. n>1000 for each plot. Scale bar: 50 μm.

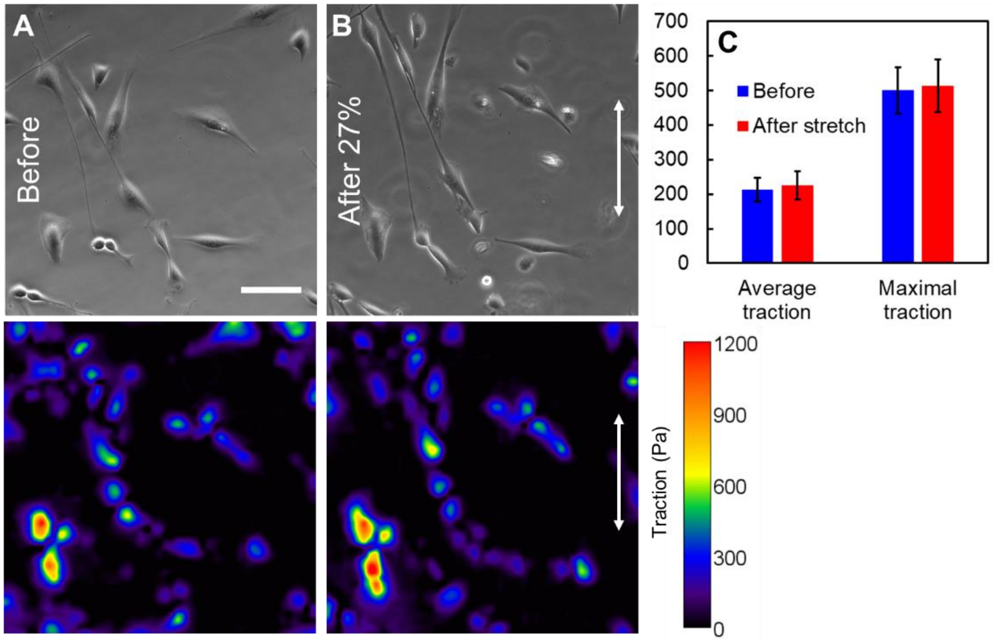


**Figure S10**. Effect of uniaxial mechanical stretch on cell traction forces. **A**. Bright field image (top) and map of traction force (bottom) for cells before the stretch. **B**. Bright field image and map of traction force for cells after a 27% uniaxial stretch in the y direction (indicated by the white arrow). The polyacrylamide hydrogel substrate is 21 kPa. Data are obtained 40 minutes after the stretch. **C**. The average and maximal traction forces of the cells before and after the stretch. n=10 for each column. No statistical differences are found (p=0.83 for average traction, p=0.91 for maximal traction). Scale bar: 100 μm.

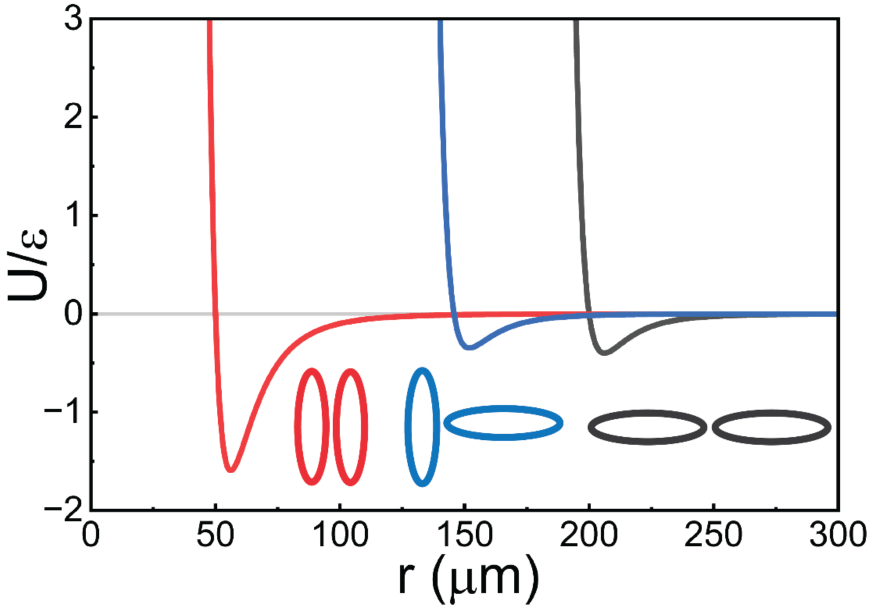


**Figure S11**. Cell-cell interaction given by a Gay-Berne potential. The potential is weakly attractive at long distances and strongly repulsive at short distances. For ellipsoid agents, the side-by-side parallel alignment is more energy favorable (red) than tiled (blue) or end-to-end (grey) configuration.

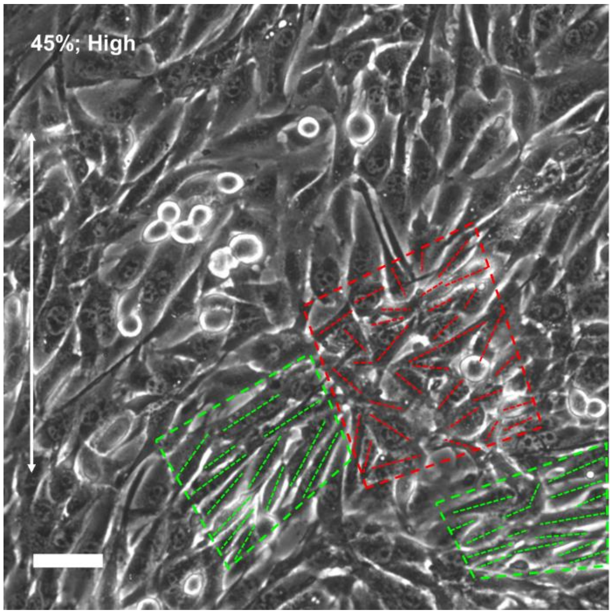


**Figure S12**. Representative area of incomplete alignment due to kinetically trapped domains in a C2C12 monolayer 24 h after cumulative 45% stretch at high density (initially 725 cells/mm²). The dashed red box marks a defect-rich, locally unaligned region that resists reorientation due to packing-induced kinetic barriers. Dashed green boxes mark misaligned parallel arrays. Short lines indicate each cell's orientation. Scale bar: 50 μm.

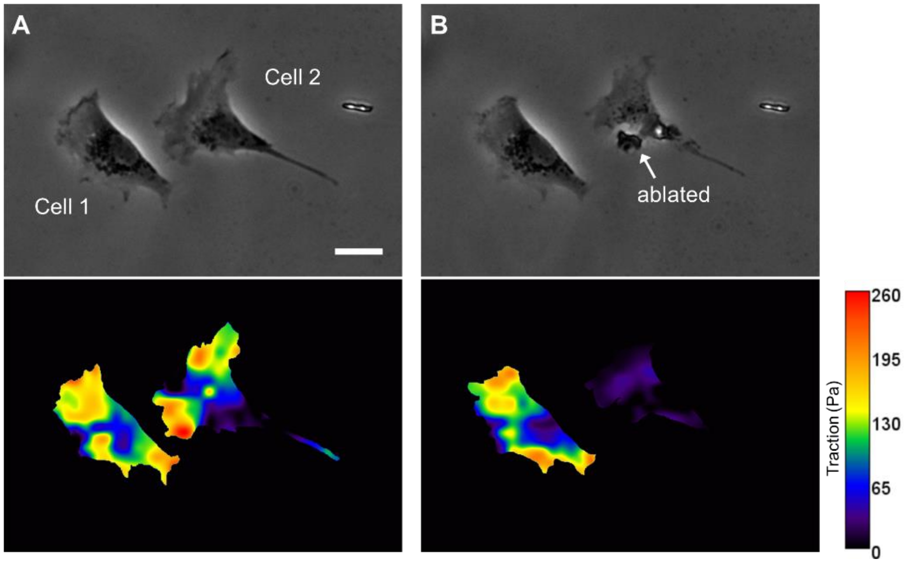


**Figure S13**. Combined laser ablation of cells and TFM to test the cell-cell mechanical interactions through the deformable substrate. **A**. Phase contrast image of a pair of cells in close vicinity before laser ablation and the corresponding traction force map. **B**. Phase contrast image of a pair of cells in close vicinity after the ablation of cell 2 and the corresponding traction force map. The polyacrylamide hydrogel substrate is 2.6 kPa. Scale bar: 50 μm.

**Supplementary Videos 1-4**. Agent-based simulations on cell alignment evolution with different cell density and configurations, corresponding to Fig. 5. Each timestep corresponds to 3 min
